# A High Dose Tango Intervention for People with Parkinson’s disease (PwPD)

**DOI:** 10.1101/613661

**Authors:** Débora B. Rabinovich, Nélida Garreto, Tomoko Arakaki, Joseph FX DeSouza

**Affiliations:** Centre for Vision Research, Department of Psychology, York University, Canada; Movement Disorders Department, Ramos Mejia Hospital, University of Buenos Aires, Buenos Aires, Argentina; Department of Biology, Neuroscience Graduate Diploma Program, Interdisciplinary Graduate Studies, York University, Canadian Action and Perception Network (CAPnet), Toronto, Canada, M3P 1P3

**Keywords:** Parkinson’s disease, rehabilitation, dance, high dose, psychological needs, partner dance, sleep

## Abstract

**Background:** Dance has been used extensively to help supplement ongoing therapies for people with PD, most commonly on a weekly or biweekly basis. A daily dose, however, may provide additional benefits. This study examines the dose effect of a dance intervention delivered within a clinic for movement disorders in which PwPD are paired with experienced studio tango dancers.

**Objective:** The current study aims to examine the dose effects of daily dance for PwPD on motor and non-motor functions directly within a movement disorders clinic.

**Design:** within-subject, pre-post-intervention, mixed-methods evaluation including UPDRS III.

**Setting:** The intervention was held at the Movement Disorders Department of a General Hospital in Buenos Aires, Argentina over two-weeks.

**Subjects:** The class had 21 people in total attendance per class. Two were expert tango dancers and instructors, nine were advanced tango dancers (volunteers), two were caregivers and eight were people with mild and moderate severity [Hoehn and Yahr (H&Y) scale 1-2] idiopathic Parkinson’s disease.

**Intervention:** Ten dance lessons, each 90-minutes daily within a two-week period.

**Outcome measures:** The Movement Disorders Society Unified Parkinson’s Disease Rating Scale (UPDRS) part III was used for pre- and post-motor assessment. Psychological questionnaires, a Likert scale examining symptoms, and a pictorial scale were used to assess non-motor aspects. Semi-structured interviews were conducted to assess the impact of the dance intervention on participants’ experience.

**Results:** Our study found a significant 18% amelioration in motor symptoms as measured by UPDRS III. We also found improvements on activities of daily living (ADL), sleep, Psychological Needs variables - post high dose dance intervention in Likert Scale.

**Conclusions:** A high dose short-term tango intervention for PwPD improved motor and non-motor aspects of PD such as ADL and sleep with high levels of adherence (97.5%) and enjoyment reported by participants. This dance intervention also improved participant’s perception of their own skills. The frequency or dosage of dance in rehabilitation suggests that an increased dose from once per week to 5 times per week can ameliorate many symptoms of PD and could be used as a short-term intervention.

## Introduction

Parkinson’s disease (PD) is a progressive neurodegenerative disorder caused by loss of dopaminergic neurons within the basal ganglia^1^. It affects nearly 1 million people in the US^1^ and five million worldwide^2^. PD is considered a movement disorder, symptoms of which include tremors, rigidity of muscles, stiffness, slowness, and impaired balance^1^. As the disorder progresses, non-motor symptoms such as cognitive impairments, slow speech, digestive problems, and sleeping disorders are observed^3,4^. In fact, Rapid Eye Movement (REM) behavioural disorder – a sleep pathology – is characterized by vivid dreams or nightmares accompanied by strong motor movements; this may precede an actual diagnosis, and is currently being examined as a biomarker for PD. Additional factors including sleep apnea, muscle rigidity, restless leg syndrome, tremor, the effects of PD medication and emotional problems exacerbate sleep difficulties^50^, leading to disruption of intrinsic circadian rhythms (sleep-wake cycles) and producing chronic issues of fatigue, irritability and impaired cognition^50^. Consequently, there is a high occurrence of emotional dysregulation observed in patients, which is aggravated from personal and social life impairments imposed by sleep disturbances. Currently, there is no cure for PD and although pharmacological interventions and other treatments temporarily reduce motor symptoms, significant impairment in motor and non-motor functioning continue to persist which, as the disorder progresses, compromises quality of life^4,8^. Rehabilitation programs based on physical activity and exercise are customarily suggested for people with PD^9^. The aim of these programs is to improve balance, strength, length of gait and range of motion^10–12^. However, traditional rehabilitation programs experience high levels of attrition^13,14^. In fact, the problem of adherence to exercise and rehabilitation programs is poorly informed^15^, but current research suggests that retention rates are particularly low for people over 60 years old^16–18^. Novel exercise interventions, including boxing, tai-chi and dancing have emerged to target the limitations presented by traditional rehabilitation therapies, enhancing social interaction and peer support^19,20^. Recently, a variety of dance interventions have gained popularity as a feasible option for PwPD^21–28^. Argentine Tango is one such intervention which has been assessed in several cohorts of PwPD^29–31^, and many studies have provided support for this intervention as a feasible tool for neurorehabilitation for PwPD^17,32^.

When comparing exercise-based programs for PwPD to those involving tai-chi or other forms of dance, Argentine Tango has been associated with equal or greater improvement in outcomes^29,33,34^. There has been limited research on the frequency or dosage of dance interventions, with only two studies completed on daily dose effect on PwPD^31,49^. Dance programs based on Argentine Tango are typically offered in a weekly or biweekly modality; however there is some precedent for offering a dance-based program with a more intensive format. Earhart and Hackney (2009) found frequent tango lessons to be an effective treatment for individuals with mild-moderate/severe PD^31^. Another study using the dance form Contact Improvisation showed improvements in balance, functional mobility and overall improvement in quality of life (Marchant, Sylvester & Earhart, 2010). Both of these studies offered 10 dance lessons in a two-week period. The benefits of increased frequency of dance rehearsal have been observed and are shown to improve muscle strength, balance, coordination, range of motion and flexibility^35^. These are the main reasons that more extensive hours of practice are suggested and used to obtain better outcomes^35^. The frequency or dosage of dance is therefore relevant in counteracting the progressive nature of this neurogenerative disorder. Improved functional outcomes for PwPD is observed through increased frequency of dance interventions, as suggested by the findings of these past studies which posit a potential link to motor training and neuroprotective effects of dance^17,36,37^.

### Enjoyment, Vitality and Psychological Needs

The future goal of this study is to assess the effects of tango class as a non-pharmacological intervention which may significantly help regulate internal rhythms, coordination, balance and mood, contributing to participants personal and social resources which may momentarily relieve symptoms of PD. The multimodal characteristics of dancing^17,38,39^ amalgamate exercise and cognitive activity with emotional and social engagement. As a result of this, the benefits of dance can be observed after only a single session of a dance intervention^40–42^. Additionally, previous qualitative research^43^ suggests that these the multimodal characteristic of dance fosters an environment where the basic psychological needs of autonomy, competence and relatedness as proposed in self-determination theories can be nurtured^44,45^. On the basis of self-determination theory (Ryan and Deci^45^), we hypothesize that tango classes will foster or nurture basic psychological needs that are the basis of the elevated sense of vitality and enjoyment most participants in dance based rehabilitation programs encounter^46^. In fact, the sense of vitality is one of the high predictors of an elevated perceived quality of life, and therapies based on dance seem to promote an increased sense of vitality^47^. Thus, this study will also aim to assess changes in psychological needs as a function of tango classes.

#### Characteristics of Partner Dancing

Few dance programs have assessed the characteristics of partner dancing and its role in dance with PwPD. Although several programs have used partner dancing, little has been explored about the specificities involved in this form. In a previous study^1^ in which we hypothesized about the underpinning mechanisms concerning dancing with an expert dancer, patients reported having experienced great pleasure, an embodied sense of security, freedom and a greater sense of mastery. In all cases, patients expressed the pleasure and comfort they experienced when dancing with an experienced dancer, and commented on how this has profoundly affected their healing process.

The main objectives of this study is to investigate the effects of a daily high dose short duration Argentine tango dance intervention for PwPD. The study has two aims. The first aim is to examine changes in motor performance, improvements in psychological needs - defined as competence, autonomy and relatedness - and self reported changes in sleep, mood and vitality. The second aim is to assess the difference between self perception of ones’ own capacities during partner dancing in comparison with practicing the sequence without a partner.

## Methods

### Participants

The class was attended by 21 people in total: 2 expert tango dancers and instructors, 9 advanced tango dancers, 2 caregivers, and 8 patients (Hoehn and Yahr (H&Y) scale 1-2) with idiopathic Parkinson’s disease who participated daily in the Argentine tango class program. The 8 people diagnosed with idiopathic PD were recruited from the Ramos Mejia Movement Disorders Department. Parkinson’s diagnostic criteria followed the UK Parkinson’s Society brain bank guidelines of bradykinesia, muscular rigidity, rest tremor, and postural instability (UK, Brain Bank Diagnostic Criteria). All patients were on Levadopa (L-dopa) medication. They took the medication one to two hours prior to the start of dance classes. Each participant demonstrated clear benefits from PD medications. 6 out of the 8 subjects had participated in previous tango for PD classes at the same hospital offered in a weekly modality during the previous year for a period of approximately 8 months. Two of them were new to the program. All were taking tango classes on a daily basis for the first time at the Movement Disorders Department of a General Hospital in Buenos Aires, Argentina (Hospital General de Agudos Jose Maria Ramos Mejia, Ciudad Autonoma de Buenos Aires, Division de Neurologia). The classes which ran within a 15-day period were taught by two experienced tango instructors (1 male and 1 female) who led all dance sessions. Each class counted on the participation of non-PD volunteers, with most being in a tango teaching program (7), while others were relatives or caregivers (2).

### Intervention

The intervention consisted of ten 90-minute classes held over a duration of 2 weeks at the Abnormal Disorders section within the Neurological department of a General Hospital in Buenos Aires, Argentina. PwPD always danced with a partner without PD, most often with an experienced tango dancer. Every session began with a 10 to 15-minute warm-up consisting of breathing, postural alignment, and body awareness exercises performed in a circle, to awaken the body and mind prior to the dancing. Classes consisted of an adapted tango class, in this case very similar to regular tango classes where new steps are introduced and some rehearsed in a new or previously presented fashion. Each gender danced in the traditional gender roles of male as leaders and females followers. However, several non-gendered exercises were introduced to practice balance, sensorial exercises, or movements associated with walking backwards (Table 1). Partners rotated during the class to allow all participants to interact with different volunteers and tango teachers. Classes were held in a 6 × 4-meter room, which usually functions as a medical gathering or conference room. Participants brought food to share during the 10-minute rest, halfway into the dance class. The hospital provided beverages such as water, tea and coffee at every class.

**Table 1:**
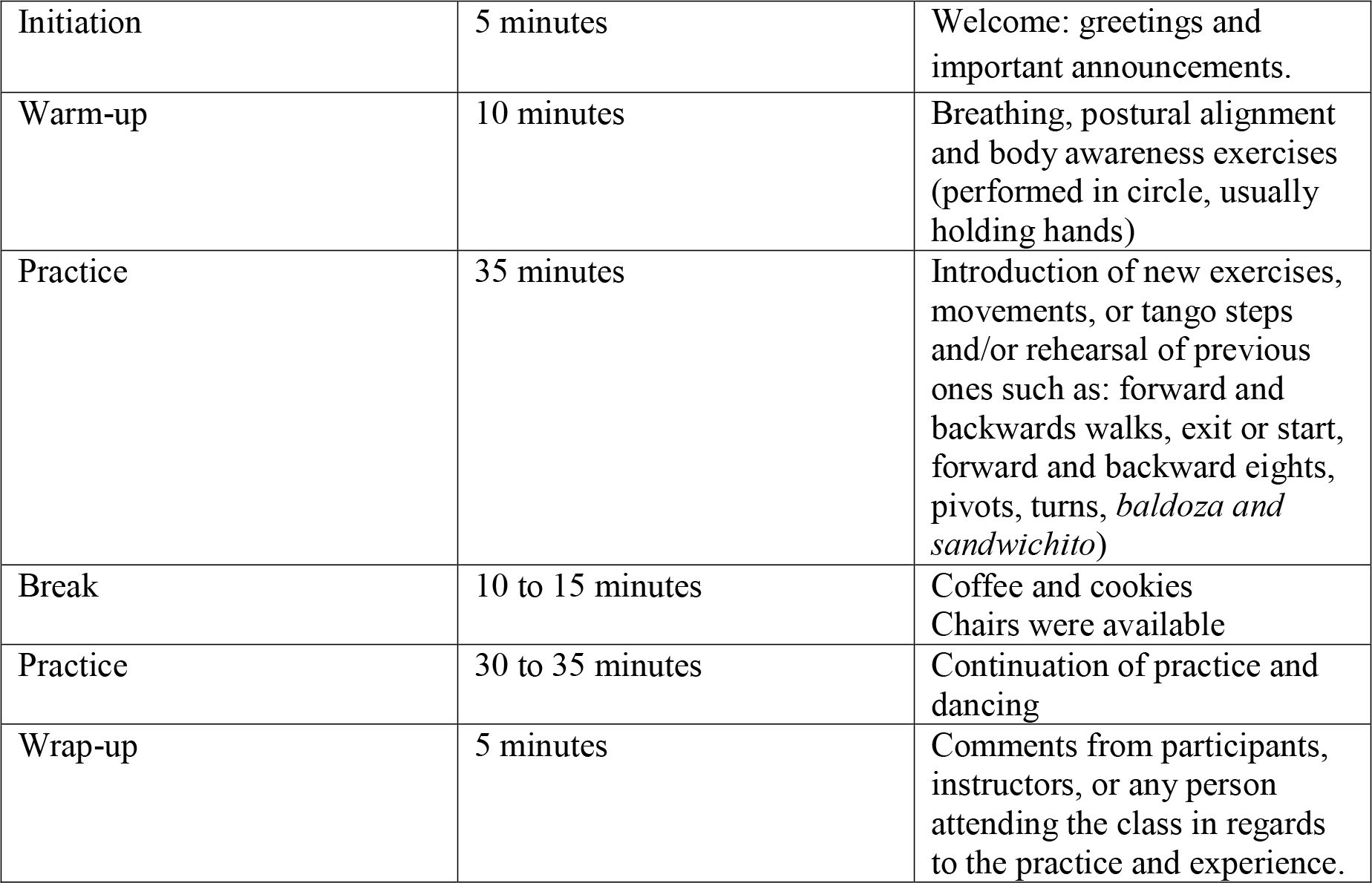
Organization of a typical 90-minute Argentine tango class

### Data Collection

The Movement Disorders Society Unified Parkinson’s Disease Rating Scale (UPDRS) part III ^52^ was used for data collection of motor symptoms pre- and post- dance intervention. A 15-item Likert Questionnaire Scale assessed perceived improvements in motor and non-motor aspects. The scale offered a 0 to 100% range to assess participant’s perception with regard to improvements in each area following the dance intervention. Positive and Negative Affect Scale (PANAS; measures mood), a Vitality Scale (7 items), a Psychological Needs Scale (3 items), and a Beck Depression Inventory were used for assessment of non-motor aspects^53,54,55^. All questionnaires and scales were validated forms of translated Spanish versions. Assessments of participants were conducted the week prior to initiation of the dance sessions and the week following completion of 10 dance sessions. Participants were tested when ON-medication at the same time of day for both pre- and post-measures. A neurologist from the neurological department of the Ramos Mejia Hospital conducted both sets of UPDRS assessments. Likert scales were completed at the Hospital the week following completion of classes and were self-administered. The scale included items that represent important concerns for this population such as mobility, ease of walking, and performance of ADL as well as mood, sleep quality, and psychological variables related to autonomy, confidence, and relatedness (basic psychological needs).

The impact of dancing on the subjective experience of participants were explored using individual semi-structured interviews. A pictorial test consisting of a sequence of 5 pictures showing different degrees of dance abilities was used as a pilot project to assess PwPD’s self-perception of their ability to dance. A similar design of 5 sequential pictures was used to measure perceived stage of Parkinson’s symptoms at their current or present moment. Both tests were assessed with participants prior to the study to validate that the pictorial test graphics clearly represented a scale of dance abilities and different stages in Parkinson’s symptoms.

### Design

This study consisted of a within-subjects design, pre- and post- longitudinal intervention and using mixed-methods evaluation. This project is the initial phase of a planned 4-year intervention.

### Ethics

This study was approved by the Ramos Mejia Ethical Committee and the York University IRB (*e2013-313*). The conduct of this investigation conformed to ethical and humane principles of research. All participants provided written informed consent prior to starting the study.

### Analysis

Changes in motor symptoms were assessed using the UPDRS pre and post scores and a student’s paired *t*-test was used at the significance level of (*p* < 0.05) for a one-tailed test -since the literature has shown that daily dance reduces motor symptoms.

## Results

All participants completed the two-week intervention consisting of 90-minutes daily Argentine tango classes. None of them dropped out of the program and only two of them missed one class (2/80 total scores of the group), each for personal activities not related to PD or the program. In both cases, they let researchers and tango teachers know about their absence in advance and regretted missing the class. Attrition rates were as follows: 6 participants attended all the classes, 2 participants attended 90% of classes; attendance rate was 97.5%.

### Interview/Questionnaires

None of the participants reported being tired or unable to complete the program even though half the participants commuted 60-90 minutes each way on the public transportation to attend classes. On the contrary, they felt very enthusiastic and energetic as the program progressed. They all reported improvements such as better quality of walking, ability to move, sleep, energy and mood. They also expressed how energized they felt by meeting with the group every day. None of them reported negative outcomes or side effects, although some reported their concerns with regards to the moment that the activity would stop, particularly how it will affect their daily enthusiasm, motivation, mood and exercise levels.

### Motor Scales

Participants significantly improved on the Unified Parkinson Disease Rating Scale Motor Subscale III, with significant paired t-test, p<0.05 (figure 1).

**Figure 1:**
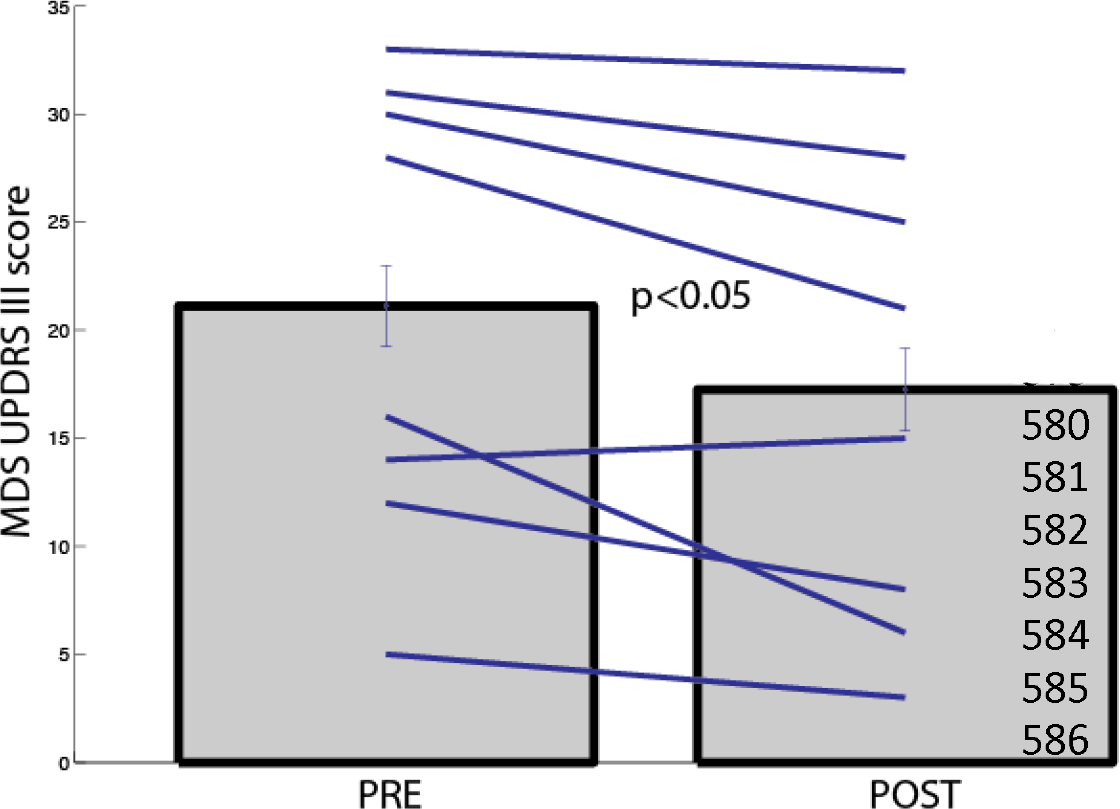
MDS-UPDRS scores for the pre vs. post times. 78 dance classes for the 8 participants between the pre/post sessions over 12 days. Error bars signify S.E.M. Each line represents one participant’s data for the pre- and post-class UPDRS scores.

### Likert Scale

The Likert scale was administered four days after completion of the last class. The Scale includes 15 items concerning general mobility, tremors, ADL, mood, relatedness, competence, and vitality. An item concerning sleep was included due to participants’ statements during interviews regarding improved quality or amount of sleep. Group Likert Scale results show the highest improvements in relatedness (Item 7 in Figure 2 = 97.5) followed by confidence in walking (item 2 in Figure 2 = 93.7) and feeling more physically active (item 10 in Figure 2 = 93.7). Participants also reported important improvements in mood, ADL, vitality and sleep.

**Figure 2:**
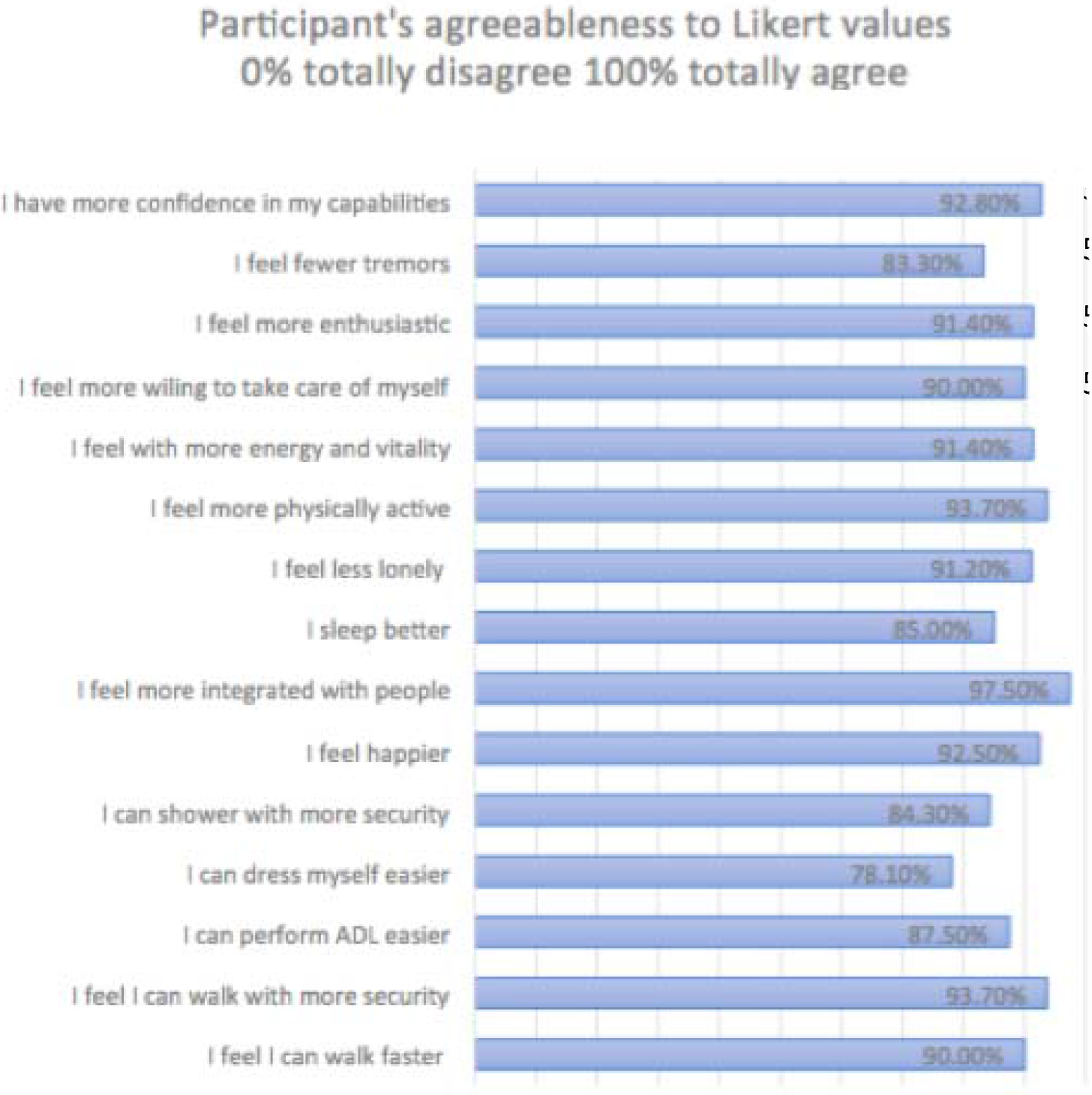
Likert scores for the post time which had the 78 dance classes for the 8 participants between the pre-/post sessions over 12 days. Each line represents the PwPD group’s data for Likert questionnaire as percentages.

#### Psychological Questionnaires

In addition, the following standard scales were administered: Beck Inventory (BD-II Spanish Version)^56^, PANAS (Spanish Version)^53^, Psychological needs^55^, and Vitality scales^54^. Since the sample was very small (n=8), and two participant’s responses were not completed correctly, no statistical analysis was performed. However, trends in patient’s responses were observed and are presented in this paper for future research.

##### PANAS

All participants completed the mood scale questionnaire prior to the intervention. Only six participants completed the post-intervention questionnaire. There was a moderate improvement in positive mood after completion of the program as well as a decrease in negative mood in four patients. One patient showed equal results for both pre and post measures, and one showed a very small increase in negative mood (1-point increase).

##### Vitality

Five patients showed improvements in vitality scores after completion of the program. One patient showed no differences between pre and post scores.

##### Psychological needs

Five patients showed slight improvements in Psychological Needs scores. Most items were in the autonomy area, followed by competence, and relatedness. Only one patient showed a slight decrease in relatedness (1 point).

##### Beck Inventory

Only six participants completed the baseline Beck Inventory test. Only two of the six completed the post intervention Beck Inventory, therefore a comparison assessment was not feasible. Five of the Beck Inventory’s baseline questionnaires fall over normal standard rates, presumably due to the moment and circumstances of the administration. These five questionnaires were completed a half hour prior to the first class.

##### Partner dance

As a pilot project, assessments for Psychological Needs, Vitality, and self-perceived competences were measured after a 10-minute practice in solo dancing compared with 10-minute partner dancing (which was always performed with a more skilled person). We used the same 3-item Psychological Needs Scale and 7-item Vitality Scale for pre and post intervention which asked, “how did you feel about this during the last two weeks”. With the new pilot project, the basic question was “how you feel at this moment”, and we compared the two different moments (after 10-minute solo practice VS 10 minutes of expert partner practice. In addition, we used a visual scale (previously presented to patients to verify its internal validity) to measure their perceived competence as dancers as well as their perceived level of PD stage. Four patients showed increases in self-perceived dance skills after completing partner rehearsal in comparison with a solo practice, while four participants maintained equal levels in both, measured by a pictorial scale (p<0.05). No significant changes were reported in the verbal scales assessing vitality and psychological needs following solo or partner dance practice.

## Discussion

This study aimed to assess the impact and potential of dance in rehabilitation for PwPD in high doses. We assessed the effects of an intensive short-term intervention using Argentine tango on motor and non-motor symptoms. As expected, our results showed significant improvements on MDS-UPDRS III scale. On average, these improvements represent an 18% decrease in PD symptoms on this scale. These results are distributed as follows: 4 patients showed major improvements between 30% - 50% (2 of whom had not participated in previous dance programs for neurorehabilitation), 2 patients showed good improvements between 18%-30%; and the remaining were the same (1 patient showed only a 1 point decrease and 1 patient showed a 1 point increase). Reasons for this distribution are not clear, but overall results indicate that on average patients largely benefited from the rehabilitation program, suggesting that intensity level or high dose interventions are a critical area for future research.

The Likert Scale aimed to assess perceived changes that would not easily be contemplated by standardized scales. Results from PwPD’s agreeableness on Likert scale questionnaire showed agreement with improvements in their ability to walk, perform ADL, sleep as well as be in a more positive mood. There was also significant improvement of psychological needs such as relatedness, competence and autonomy. These results suggest that the dance intervention is perceived as a very enjoyable activity, which is in line with previous qualitative research findings. Several authors have proposed that activities which foster an environment where these basic psychological needs are nurtured promotes a greater sense of vitality and enjoyment. The perception of the dance intervention as an enjoyable activity comes from the greater sense of autonomy, relatedness and competence that PwPD experience, which is in line with previous qualitative findings. Aside from this augmentation on the Psychological Needs variables (Items 7,9,11,15 on Likert scale), rates in the 3-item Psychological Needs Scale did not improve as expected when comparing baseline with post intervention scores. These results may correlate to the high scores shown in the pre-intervention test which leaves very little space for improvement. Reasons for these high baseline levels are not clear but may be related to the fact that most participants had already participated in a weekly dance intervention.

### Beck Depression Inventory (BD-II)

This study included Beck Depression Inventory since it is a common measure to assess depression. Again, high baseline scores left little room for improvement. In addition, 3 participants failed to complete one of the two scales (pre and post intervention) making analysis more difficult.

### Semi-structured interviews

During the semi-structured interviews participants expressed the importance of the group and camaraderie, the relevance of dancing with the teachers and with skilled dancers, and the importance of focusing on their movement, balance, and abilities instead of exclusively focusing on the things they were not able to do. They expressed the importance of the classes strengthening their motivation and providing a sense of challenge for attending the program daily. They felt fortunate to have the chance to participate in the program and the fact that dancing helped them to better cope with the disease and associated symptoms.

#### Limitations

This longitudinal study had a small sample size (n=8). There was no control group and the instructors all knew who the people with PD were. All participants had the desire to take dance classes and this could be a bias for the success rate or improved vitality or enjoyment of the activity. Another limitation of this study was the self-administered questionnaires with most baseline tests administered in a group setting. The Likert questionnaire is a self report and as such, is subject to the inherent biases in all self report questionnaires. These questions are framed such that they suggest that the patients should feel better after the intervention and this reduces the validity of the questionnaires. Also, the five questionnaires such as BD-II, vitality and psychological needs questionnaires amongst others, were completed a half hour prior to the first class and when the participants were thus excited. Additionally, there was not enough privacy for each of them in this room, which may have biased their answers, thus potentially affecting the results. High improvements in sleep quality and quantity were not one of the expected results of the program so no specific tests or assessments other than the Likert Scale questionnaire was included. However, previous studies found improvements in sleep following a tango intervention for people with self-referred affective disorders, and it may be similar mechanisms involved in our PD intervention. Further research should address sleep as a relevant non-motor aspect that may contribute to a better understanding of the extent of benefits associated with high dose dance classes for PwPD.

## Conclusions

Daily dance lessons completed within a short period appear to be appropriate and effective for individuals with mild-to-moderate (I and II, H&Y) Parkinson’s disease. Patients showed significant improvements in motor symptoms measured by UPDRS part III. In addition, a Likert Scale completed after finalization of program showed improvements in perceived mobility, ability to walk, and performance of ADL as well as better quality and quantity of sleep and improved mood. It also shows that the activity fosters Psychological Needs such as competence, relatedness and autonomy. This study also aimed to investigate the nuances of partner dancing. PwPD showed improvements in their self-perceived skills after rehearsing with a more experienced dancer in comparison with a solo dance practice. This is in line with former research and deserves further examination to understand the underpinning neural mechanisms. As in any other treatment program, the question of the intensity level or dosage deserves deeper examination to quantitatively determine the peak potential of the intervention. When performed at high intensity level, the two-week Argentine tango classes for PwPD showed significant improvements in motor symptoms and no side effects. The results from this study correspond with former research and deserve further exploration. High dose intervention proved to be beneficial for people who had previously attended a weekly tango class as well as those who had never taken classes before. As previously stated, dance is a non-pharmacological approach that helps maintain health through physical activity and cognitive training, offering an enriched environment in which to socialize and learn new skills. As a form of rehabilitation in PD our findings present hope for the possibility of decreasing the rate of progression of the disease.

## Acknowledgements

We would like to thank Juan Manuel Firmani and Veronica Litvak for their dedication and involvement in teaching and facilitating this program. We would also like to acknowledge the volunteers for their participation and strong commitment to the program. T. Chosang, R. Cohan, K. Bearss commented on manuscript. Tenzin Chosang also assisted with the final editing of the manuscript.

## Footnotes

1 Idem 1

